# Behavioral responses of the invasive fly *Philornis downsi* to stimuli from bacteria and yeast in the laboratory and the field in the Galapagos Islands

**DOI:** 10.1101/696492

**Authors:** Boaz Yuval, Paola Lahuatte, Arul J. Polpass, Charlotte Causton, Edouard Jurkevitch, Nikolaus Kouloussis, Michael Ben-Yosef

**Author notes:** Address correspondence to B. Yuval.

## Abstract

*Philornis downsi* (Diptera: Muscidae) is a nest parasitic fly that has invaded the Galapagos archipelago and exerts an onerous burden on populations of endemic land birds. As part of an ongoing effort to develop tools for the integrated management of this fly, our objective was to determine its long and short-range responses to bacterial and yeast cues associated with adult *P. downsi*. We hypothesized that the bacterial and yeast communities will elicit attraction at distance through volatiles, and appetitive responses upon contact. Accordingly, we amplified bacteria from guts of adult field-caught individuals and bird feces, and yeasts from fermenting papaya juice (a known attractant of *P. downsi*), on selective growth media, and assayed the response of flies to these microbes or their exudates. In the field, we baited traps with bacteria or yeast and monitored adult fly attraction. In the laboratory, we used the Proboscis Extension Response (PER) to determine the sensitivity of males and females to tarsal contact with bacteria or yeast. Long range trapping efforts yielded two female flies over 112 trap nights (one in extracts from bird faeces and one in extracts from gut bacteria from adult flies). In the laboratory, tarsal contact with bacterial stimuli from gut bacteria from adult flies elicited significantly more responses than did yeast stimuli. We discuss the significance of these findings in context with other studies in the field and identify targets for future work.

---

*Philornis downsi* (Diptera: Muscidae) is a nest parasitic fly that invaded the Galapagos archipelago in the mid twentieth century (Causton et al., 2006; Fessl et al., 2018; Kleindorfer & Sulloway, 2016). Female flies oviposit in active bird nests. Post hatching, larvae undergo three instars, during which they typically feed on the nestlings present in the nest, first by acquiring blood from vessels in the host’s nares (during the first instar), and subsequently by nocturnal feeding bouts on blood and tissues of the host (Kleindorfer & Dudaniec 2016; McNew & Clayton 2018).

The broad host range of these flies, lack of competitors and natural enemies, their high dispersal ability and adaptability to harsh environments have all contributed to their successful invasion (Fessl et al., 2018). This success is manifest in the impact on the local passerines. Since *P. downsi* was first observed in 1997 (Fessl et al 2001), nearly all of the passerines on the islands have been recorded as hosts (Fessl et al., 2018). Furthermore, the intensity of parasitism (number of larvae per nest), has been rising, with attendant increase in mortality of the defenseless hosts. Particularly susceptible are species of the endemic (and iconic) group of birds known as “Darwin’s finches” (Kleindorfer & Dudaniec, 2016; Fessl et al., 2018). Thus, for example, the Medium Tree-finch (*Camarhynchus pauper*) has lost >50% of its population on Floreana island (e.g., Dvorak et al.,2017). Elsewhere, on the island of Santa Cruz, populations of the Warbler Finch (*Certhidea olivacea*) are declining as levels of parasite infestation continue to rise (Cimadom et al., 2014). On the island of Isabela, the last population of the critically endangered Mangrove Finch (*Camarhynchus heliobates*) is literally on the verge of extinction, due to the combined effects of predation, habitat loss and *Philornis* parasitism (Fessl et al., 2010).

Due to the protected status of the Galapagos Islands, a control approach based on indiscriminate application of insecticides (which may be effective) is out of the question. Accordingly, an international consortium of researchers, coordinated by the Charles Darwin Foundation (CDF) and the Directorate of the Galapagos National Park in Puerto Ayora, Santa Cruz, is seeking to develop and implement strategies and tools for the management of *P. downsi* in the Galapagos Islands (Causton et al., 2013). In the short term, these approaches consist of captive breeding of nestlings of the most endangered species (Cunninghame et al., 2017), application of larvicide to nests (Knutie et al., 2013; Fessl et al., 2018), and adult trapping (e.g., Cha et al., 2016). The long-term vision is an integrated use of biological control with the sterile insect technique (SIT). Although several promising natural enemies have been identified in the Americas (e.g., Bulgarella et al., 2017), implementation will take a while as the candidate enemies need to satisfy regulatory requirements prior to introduction into the fragile ecosystem of the Galapagos Islands. Concurrently, implementation of the SIT is hampered by vast lacunae in our understanding of the basic biology of *P. downsi*, as exemplified by the extreme difficulty of rearing this fly in captivity (Lahuatte et al., 2016), our ignorance of the particulars of its mating system, and patterns of its behavior in the field.

Complementary to these approaches are methods to manipulate the microbiome of the target insect. The relationship between insects and microorganisms has received much attention in the past two decades, and hardly needs an introduction, as the contributions of symbionts to host nutrition, environmental adaptation, immunity and ultimately, fitness, are quite well known (e.g., Douglas 2015). In theory, once the specific microbial partners of an insect, and their effects on the host are identified, it is possible to manipulate them in a manner that benefits the goals of control operations (reviews by Zindel et al., 2011, Jurkevitch 2011; Yuval et al., 2013. For examples see Ben-Ami et al., 2010; Gavriel et al., 2011; Hoffman et al., 2011).

In flies (Diptera), gut and environmental bacteria have been found in many instances to provide cues that are important in foraging for food (Wong et al. 2017; Leitao-Goncalves et al., 2017; Akami et al., 2019), oviposition sites (Romero et al., 2006; Lam et al., 2007; Tomberlin et al., 2012; Zheng et al., 2013) and mates (Hoyt et al., 1971; Sharon et al., 2010). Furthermore, volatiles from bacteria and fungi have been shown to attract flies and to have potential for enhancing trap catches (Robacker et al., 1998). Indeed, Cha et al., (2016), found that *P. downsi* were attracted to traps containing active brewer’s yeast (*Saccharomyces cerevisiae*) and in particular to acetic acid and the ethanol produced by the yeast. Currently, the standard method of capturing *P. downsi* in the field is with McPhail traps baited with fermenting papaya juice. This attractant is superior to commercially available fly attractants, yet it is cumbersome and many non-target flies, wasps and moths are trapped, together with fewer than 1 *P. downsi*/trap/day (Lincango & Causton, unpublished report, CDF 2009). Clearly a selective, efficient trap would be a welcome addition to the tools employed for the study and control of this fly.

We believe that there are several possible uses for microbes in *P. downsi* management such as improving diets for mass rearing for SIT and for developing effective attractants for use in traps. Recently, we characterized the bacterial microbiome of *P. downsi* (Ben-Yosef et al., 2017). We found that larval and adult microbiomes are dominated by the phyla Proteobacteria and Firmicutes, with communities that significantly differ between life stages, reflecting the different dietary needs of the larvae and adults. In light of the widespread importance of the microbiome in shaping behavior of many insects (references above, also see Leroy et al., 2011; Wong et al., 2015), we hypothesized that the bacterial and fungal communities associated with adult *P. downsi* will elicit attraction at distance through volatiles, and appetitive responses upon contact. Accordingly, we amplified bacteria from guts of adult field-caught individuals, and yeast from fermenting papaya juice, on selective growth media, and assayed the response of flies to these microbes or their exudates. In the field, we baited traps with bacteria or yeast and monitored adult fly attraction. In the laboratory, we used the Proboscis Extension Response (PER) (e.g., Yuval & Galun, 1987; Wong et al., 2017) to determine the sensitivity of males and females to tarsal contact with bacteria or yeast.

## Materials and Methods

Traps and Baits – We devised a modified McPhail trap to hold a 10 cm. Petri dish inoculated with bacteria or yeast (Figure 1). We deployed these traps on two locations on the Island of Santa Cruz: El Barranco (−0.738117; −90.301656), a lowland arid site and Los Gemelos (−0.626379; −90.380697), a highland, humid site (Jackson, 1993). These are areas where *P. downsi* is commonly collected. Traps were deployed overnight on 8,9,10,14,15 February and 7,8,9 March of 2018, and on 11,13,15 February 2019, for a total of 112 trap nights. These periods fall within the nesting season of the local finch species (Grant, 1986). Traps were placed ∼3-5 meters high on branches of trees and spaced at least 10 meters apart.

**Figure 1.**
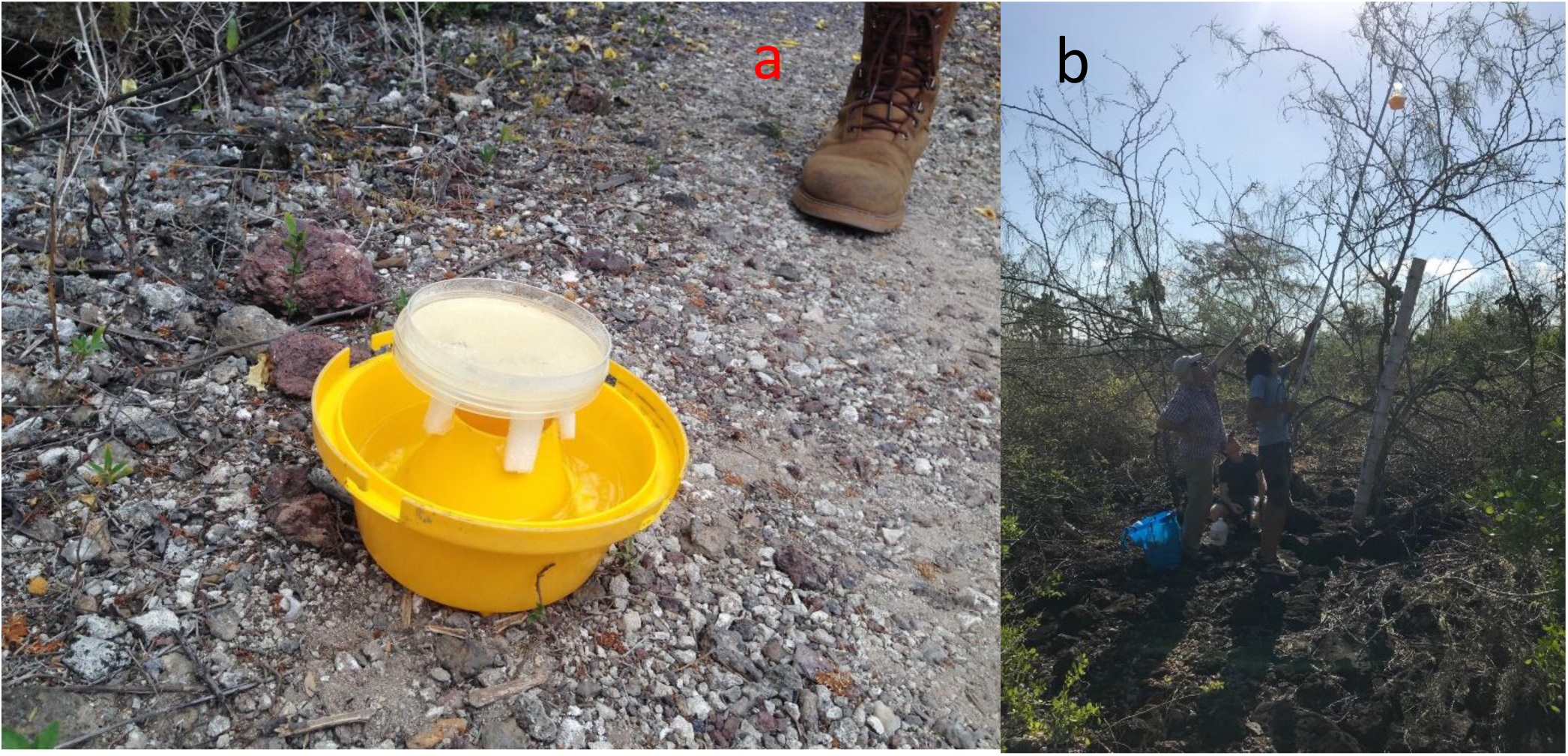
a. Modified Mcphail trap with bacterial bait (trap cover not shown). b. Positioning trap in El Barranco, Santa Cruz Island, Galapagos.

To assay attraction to bacterial volatiles, we dissected the gut of freshly caught, surface sterilized male and female *P. downsi*, and homogenized the gut contents in Phosphate Buffered saline (PBS). Homogenates were pooled (n= 3-5), and inoculated on plates with one of Luria-Bertani (LB) general agar media, Brain Heart (BH) general agar media, or Tomato agar (TA) (selective for *Lactobacilli)*. Twenty-four hours following the inoculation, the growth plates, with the bacterial colonies, were positioned (uncovered) as bait in the traps (figure 1). Concurrently, bacterial samples were taken for future identification, as described in Ben-Yosef et al., (2017). In addition to bacteria from flies, we also cultured bacteria from fresh feces of a Medium Ground-Finch (*Geospiza fortis*) on LB plates and baited traps in a similar manner. To assay attraction to yeast volatiles, a sample from 4-day old fermenting papaya juice was inoculated on Yeast Peptone Dextrose (YPD) plates, and the plates were positioned in traps. Samples from the resulting yeast colonies will be sequenced for identification based on the 18S ribosomal RNA.

Traps baited with microbe-inoculated plates or sterile plates as controls, and supplemented with odor-free soapy water at their base (to immobilize insects entering the trap), were positioned between 1630-1830 hrs., and collected between 0730-0830 on the following day. At the time of collection, we determined the number and sex of *P. downsi* in each trap, and also counted the total number of other flies and any other insects present.

PER assay – The PER experiment was done on two populations of flies. The first were adults that emerged from pupae recovered from nests that were parasitized by the fly and collected after fledglings left the nest or died. The nests were brought back to the laboratory and pupae removed from the nest material. Following emergence, flies were held in sex specific cages (dimensions: 45×45×45 cm.), and provided with a diet of bacto-yeast powder (0.87 g.) added to10 ml. of 4-day old fermented papaya juice, plus ripe berries of Muyuyo (*Cordia lutea*) and water *ad lib*. On the day of the experiment they were 6-9 days old.

The second group were adult flies caught in McPhail traps baited with fermented Papaya juice placed around the CDF headquarters in Puerto Ayora (−0.741425; −90.302766), and brought back to the laboratory between 26.1.2019 and 2.2.2019, and thus were at least 14-21 days old on the day of the experiment (15.2.2019). Following capture, they were held in cages as described above, and fed on mixture of pulped papaya (34.2%), protein powder (2.6%), egg powder (3%), milk powder (2.6%), sugar (7.9%) and water (50%).

On the evening before the experiment food was removed from the fly cages, while water remained. Four hours prior to the experiment, flies were glued with liquid silicon to the base of a disposable pipette tip, and allowed to rest in a humidified environment (Figure 2).

**Figure 2.**
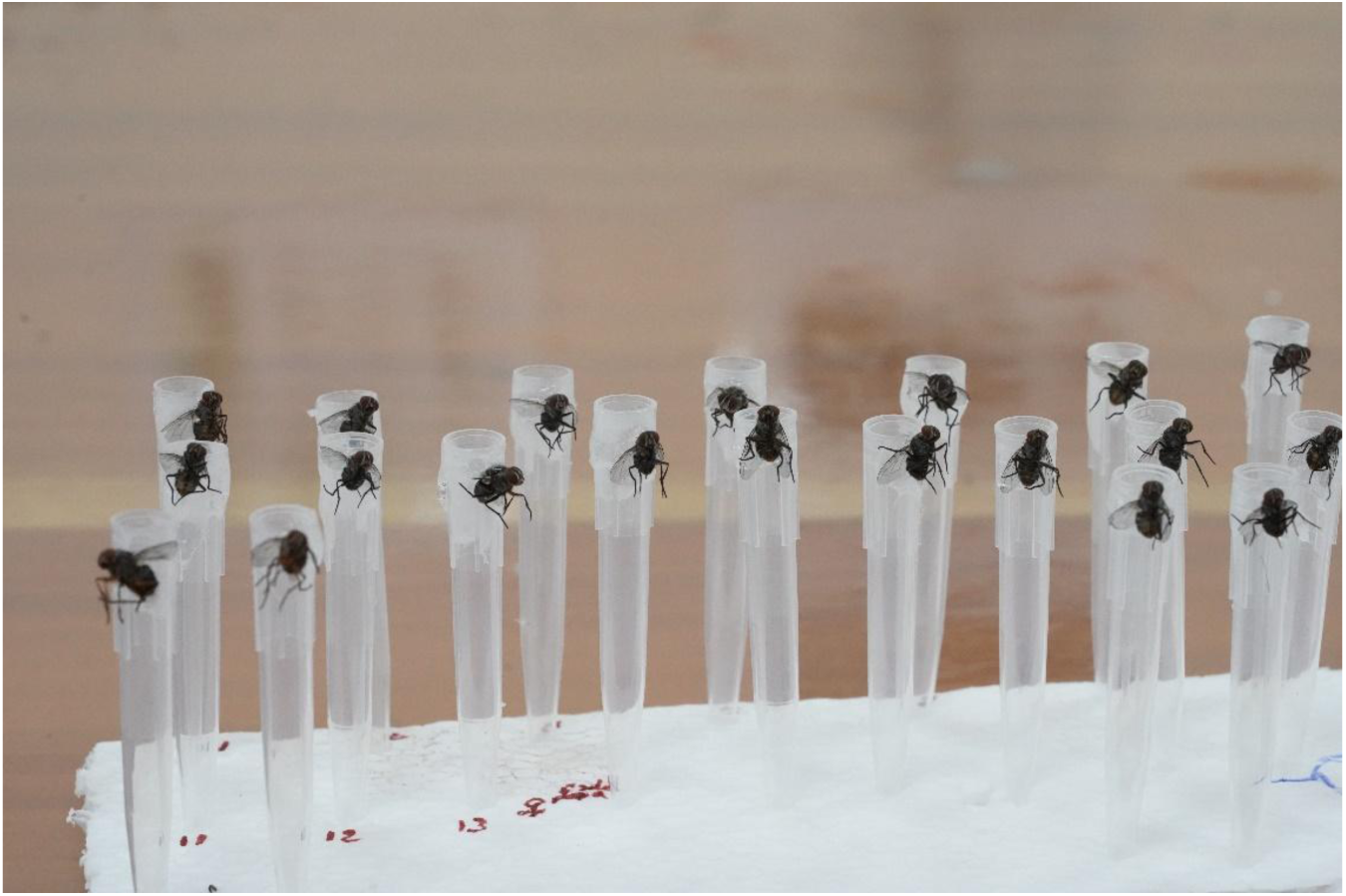
*Philornis downsi* adults prepared for the PER experiment.

The assay itself consisted of dipping the tarsi of the flies into a series of stimuli, and recording the response of the proboscis. The stimuli were presented as droplets in the following order: Water (allowing flies to drink their fill); PBS (negative control); bacteria or yeast (test stimulus); 20% sucrose (positive control), as shown in Figure 3. Flies responding to the negative control (5 females from pupae) and those ***not*** responding to the positive control (3 females and 3 males derived from pupae, and 2 trapped females) were excluded from the analysis. Full extension of the proboscis in response to tarsal contact with the test stimulus (bacteria or yeast) was scored as a positive result.

**Figure 3.**
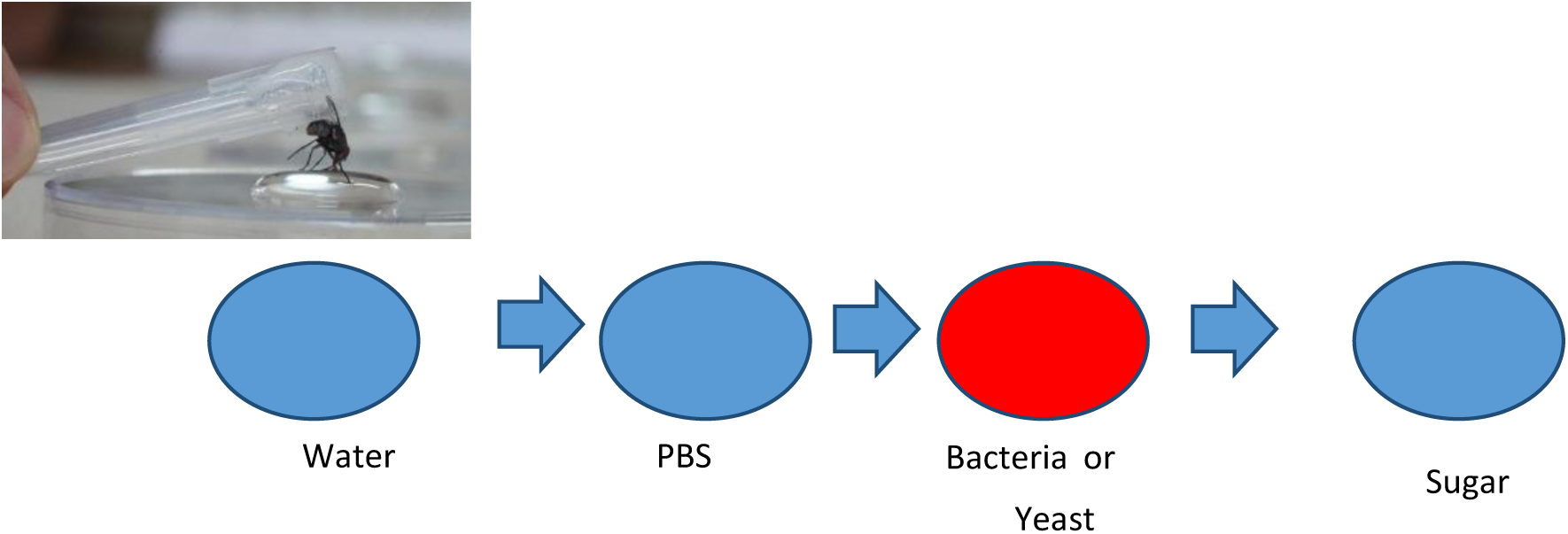
Sequence of stimuli presentation in PER experiment. Flies were allowed to drink water if they responded to it, prior to presentation of subsequent stimuli.

Bacteria were derived from the guts of three field-collected females. The guts were excised, and suspended in PBS, then plated on LB plates. After 24 hours, colonies were harvested and re-suspended in PBS. A 50 µl droplet with an estimated density of 10^5^ – 10^6^ bacterial cells was then pippeted onto a sterile microscope slide and presented as the test stimulus. For yeast derived stimuli the same approach was used. In this case yeast from fermenting papaya juice were cultured on YPD prior to harvesting and resuspension in PBS.

Statistical analysis – As can be seen below, there was no point in analyzing the trap catches. The results of the PER experiment were analyzed using generalized linear model (GLM) with Binomial distribution and log link function. JMP Pro version 14 (SAS Institute Inc.) was used.

## Results

### 1. Trapping

Over a total of 112 trap nights, we caught only 2 *P. downsi* females (Table 1). One was caught in a bacteria/LB trap, and the other in a finch faeces/LB trap. Interestingly, a variety of other flies were caught, mainly sarcophagids, calliphorids and tephritids. However, a detailed analysis of these catches is irrelevant to our goals. Also of interest was the observation that few other insects, namely wasps and moths, were attracted.

**Table 1.** Trapping results for bacterial attractants in modified McPhail traps.

| Location | No. trap nights | Bait/medium | No. of *Philornis* | Mean no. (±SD) of other flies/trap | Other insects trapped |
| --- | --- | --- | --- | --- | --- |
| El Barranco | 20 | Bacteria/ LB | 1 (female) | 4.6 (5.2) | 1 ant, 3 wasps* |
| El Barranco | 8 | Finch feces/LB | 1 (female) | 5.2 (2.7) | 8 ants |
| El Barranco | 3 | control/ LB | 0 | 5.6 (5.1) | 1 wasp |
| El Barranco | 12 | Bacteria/ TA | 0 | 14.6 (16.2) | 2 ants, 3 wasps |
| El Barranco | 3 | control/TA | 0 | 15.6 (16.8) | 1 wasp |
| El Barranco | 8 | Bacteria/BH | 0 | 3.3 (1.9) | 3 ants, 21 wasps, 3 moths |
| El Barranco | 2 | control/BH | 0 | 1 (1.4) | 2 wasps |
| El Barranco | 8 | Yeasts/YPD | 0 | 0.3 (0.7) | 1 ant, 1 wasp |
| El Barranco | 2 | control/YPD | 0 | 1.5 (2.1) | 1 ant 1 wasp |
| Los Gemelos | 16 | Bacteria/ LB | 0 | 1.8 (1.6) | 0 |
| Los Gemelos | 4 | Bacteria/ LB | 0 | 2 (2.1) | 0 |
| Los Gemelos | 16 | Bacteria/ TA | 0 | 2.8 (2.5) | 1 moth |
| Los Gemelos | 8 | Fermented Papaya/ TA | 0 | 1.8 (1.5) | 1 Cockroach |
| Los Gemelos | 2 | control/TA | 0 | 7.5 (4.9) | 0 |

\* all wasps were *Polistes versicolor*

### 2. PER experiment

The PER experimental results were very conclusive (Figure 4). The bacterial stimulus elicited significantly more responses than the yeast stimuli. The whole model analysis, which included the stimulus tested, the source of the flies (trapped or emerged from pupae collected from nests) and their sex, was marginally significant (GLM Maximum Likelihood x^2^ = 6.31, P = 0.09, n = 77). This model tested the effects individually and revealed that the stimulus (bacteria or yeast) was significant (x^2^ = 5.96, P = 0.014), while the source of the flies (x^2^ = 0.035, P = 0.85), and their sex (x^2^ = 0.06, P = 0.79), were not significant in eliciting a different response to bacteria or yeast.

**Figure 4.**
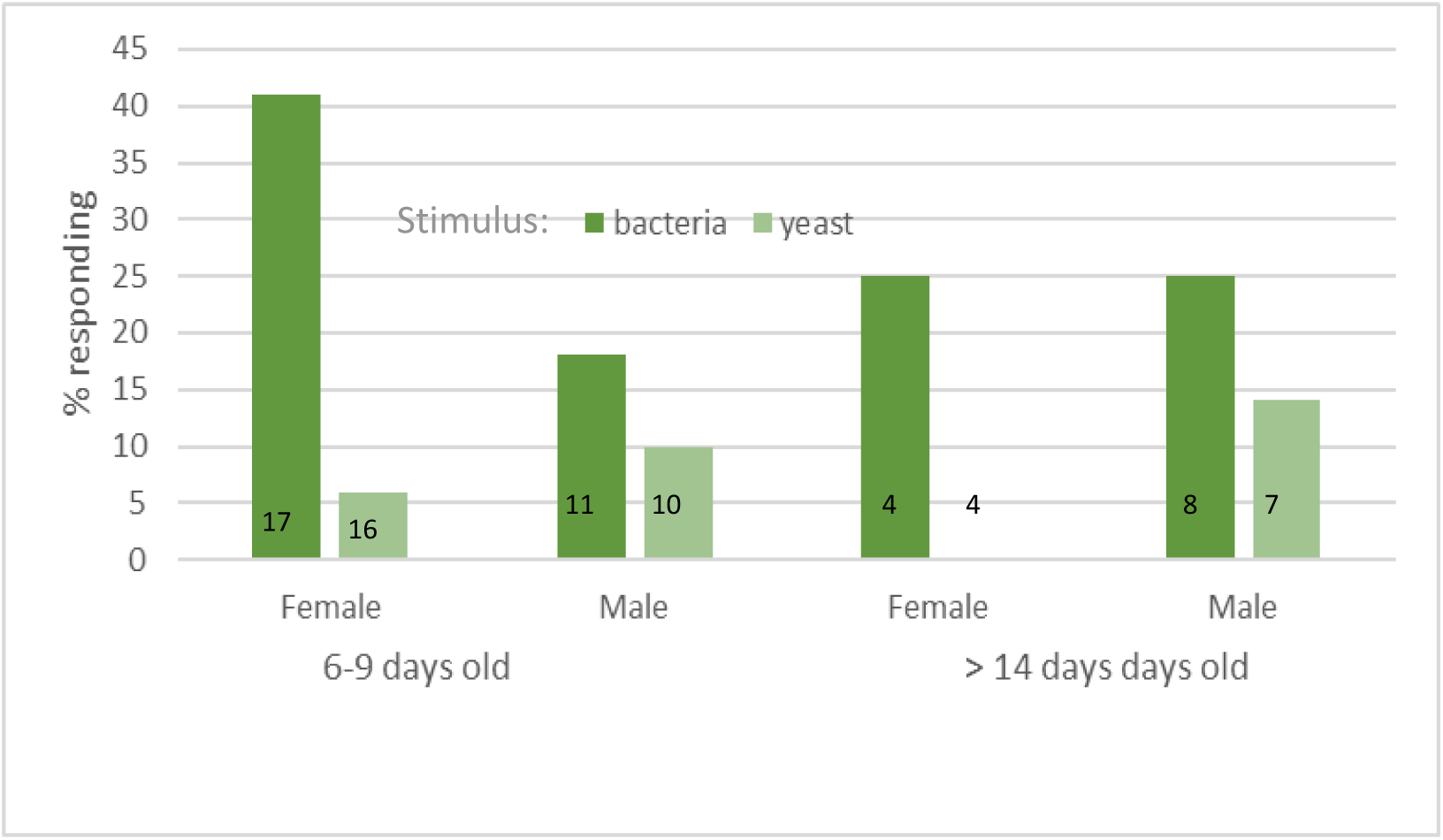
Proboscis extension responses of *P. downsi* to bacterial or yeast stimuli. 6-9 day old flies were derived from pupae recovered from naturally infested nests. Flies >14 days old were trapped in the field and maintained in the laboratory prior to testing. The difference in the response to stimuli is highly significant (GLM effect test x^2^ =5.96, P 0.014). Numbers within columns denote sample size.

On excluding the source of the flies (which showed no differences) from the analysis, the picture becomes even stronger (General Linear Model x^2^ = 6.6, P = 0.035, n= 77). While there were still no differences between the responses of males or females (x^2^ = 6.3, P = 0.51), the response to the stimuli differed greatly (x^2^ = 34.5, P < 0.0001).

## Discussion

The development of a selective and efficient trap for *P. downsi* has been identified as a research priority and would greatly enhance future integrated management programs (Causton et al., 2013). In light of published studies of attraction of other flies to microbial volatiles (see introduction), we had high hopes for our novel modified traps. Sadly, we cannot but conclude that this approach has failed. Previous studies aimed at assaying various attractants for this fly resulted in much higher catches per trap. Lincango and Causton (unpublished report, CDF 2009) review attractants that were tested between 2007–2009, among them- milk powder, Biolure, Tricocene, Muyuyo berries (*Cordia lutea*), Amines, Indole, Putrecine, Methyl-amine, and fermented papaya juice. The latter proved to be the strongest attractant, and is currently used in all monitoring efforts (Causton et al. in review). Recently Cha et al., (2016) identified a number of attractive volatiles from fermenting baker’s yeast, pinpointing acetic acid and ethanol as the most potent. The most attractive bait they assayed was a liquid combination of yeast and sugar. Extrapolating from the results of both these studies indicates that the best attractants (fermented papaya juice and active yeast), yield captures approaching (but not exceeding) 1fly/trap/day. Our results come nowhere near, and even if we consider that in 2018 our traps were deployed very early in the nesting season, this cannot be said for 2019, when nesting was in full swing and papaya-baited traps were capturing many flies. We left our traps out overnight, but they were available to the flies for at least 2 hours in the evening, and 2-3 hours in the morning, so even if they are not active in the dark, they had ample opportunity to access the baits. Indeed, on the few occasions when traps were not collected immediately, results were the same.

Interestingly, one of the females we caught was in a trap baited with bacteria from the feces of a Medium Ground-Finch (*Geospiza fortis*). The structure of the bacterial community of this sample will be detailed in the future (Polpass, in preparation). The feces we used were from a fresh deposit made by an individual frequenting the cafeteria at CDF. A recent study by Knutie et al., (2019) examined the effect of proximity to humans on the microbial community in finch feces. They found that proximity lowered the diversity of the microbiome in this species.

The other female we trapped was attracted to a community of gut bacteria from adult flies growing on LB medium. This community has been characterized in full (Ben-Yosef et al., 2017), and contains taxa such as Enterobacteriacae that in other systems are known to be important in attraction (e.g., Lam et al., 2007; Robaker et al., 1998). Our results do not immediately suggest that this microbial community magnetizes foraging flies. However, the mechanism whereby female flies localize host nests or food sources is as yet unknown, and we feel that the microbial dimension may have an important part to play. That said, our result of 2 flies in 112 trap nights are sobering (these may very well be the two most expensive flies in the history of entomology) and a change in our methodology is mandated. Nevertheless, we feel it is important to share our results, and welcome any suggestions for improving our approach.

The PER has been used widely to characterize the sensitivity of chemical receptors of flies and bees (e.g. Simcock et al., 2014; Lau et al., 2016; Mustard et al., 2019) and as a conditioned response in learning trials (e.g., Yuval & Galun 1987; Arien et al., 2015; Liu et al., 2015, 2017). In our experiment, we tested the response of adult flies to a solution containing either bacteria derived from an adult female fly, or yeast growing in fermented papaya juice. The response to sucrose at the end of the test unequivocally shows that all flies were motivated to ingest food, and the exclusion of flies responding to PBS assures us that the responses we considered were indeed specific to the test stimuli (bacteria or yeast).

We used flies from two populations in this experiment, young individuals reared from pupae, and older flies that were trapped in the field at an unknown age and maintained at length in the laboratory. Although both responded in a similar manner to the stimuli, such that it was indistinguishable statistically, the trend observed was that the older flies were less responsive overall (Figure 4). Apart from their age, these groups differed in the adult diet they received, and possibly in sexual experience. The young group were all virgins, while there is a high probability that the older group had mated prior to capture. The microbiome of flies is known to affect their responses to nutritional cues (Wong et al., 2017; Leitao-Goncalves et al., 2017; Akami et al., 2019). Different diets support different gut microbial communities, which in turn may affect behavior in a different manner. It may very well be that the different diets ingested in the laboratory (and previously in the field, by the older flies) affected the magnitude of the responses observed, if not their direction.

We found that both females and males (but especially the females) from the two populations we assayed showed a significantly higher response to the bacterial cues. Overall, 38% of females (n=21) and 21% of males (n=19) responded to bacteria, while only 5% of the females (n=20) and 11% of the males (n=17) responded to the yeast (Figure 3). Thus, although volatiles from fermented yeast and fermented papaya juice attract flies to traps, they rarely elicited proboscis response following contact with the flies’ tarsi. The elevated response to bacterial cues is intriguing. All responding flies were eager to eat, and the obvious conclusion is that the bacterial cue is associated with a food substrate. Although we do not know exactly what the main sources of nutrition are it is reasonable to assume that *P. downsi* are polyphagous and ingest decaying organic matter, fruit juices, and possibly pollen and nectar (Kleindorfer & Dudaniec 2017; Fessl et al., 2018; McNew & Clayton, 2019). While many of these substrates are associated with bacteria, they are frequently also associated with yeasts, which elicited a significantly lower response. Clearly, more work needs to be done. Specifically, to identify if the flies are responding to a specific bacterial species or metabolic product, and how the experiential and symbiotic status of the fly influences its responses. We suggest that the PER paradigm will be extremely useful in the future to answer these questions.

## Acknowledgements

We thank Alejandro Mieles for many helpful suggestions; Sabrina McNew and Courtney Pike for providing the nests; Andrea Cahuana, Ismael Ramirez, Magally Infante and Simon Heimpel for help in the field. This work was funded by a grant from the US-Israel Binational Science foundation (BSF #2016046), and by funding from Galapagos Conservancy, and from Lindblad Expeditions-National Geographic. Permission to conduct this study was granted by the Galapagos National Park Directorate (Project: PC- 18-16 & 07-18: Control of the Invasive Parasite, *Philornis downsi* and its Impact on Biodiversity) and the Ecuadorian Ministry of the Environment. This is contribution number xxx of the Charles Darwin Foundation for the Galapagos Islands.

